# Effects of dim light at night in C57BL/6J mice on recovery after spinal cord injury

**DOI:** 10.1101/2023.09.15.557980

**Authors:** John C. Aldrich, Ashley R. Scheinfeld, Sydney E. Lee, Kalina J. Dusenbery, Kathryn M. Mahach, Brigid C. Van de Veire, Laura K. Fonken, Andrew D. Gaudet

## Abstract

Spinal cord injury (SCI) can cause long-lasting locomotor deficits, pain, and mood disorders. Anatomical and functional outcomes are exacerbated by inflammation after SCI, which causes secondary damage. One promising target after SCI is manipulating the circadian system, which optimizes biology and behavior for time of day – including neuroimmune responses and mood-related behaviors. Circadian disruption after SCI is likely worsened by a disruptive hospital environment, which typically includes dim light-at-night (dLAN). Here, we hypothesized that mice subjected to SCI, then placed in dLAN, would exhibit worsened locomotor deficits, pain-like behavior, and anxiety-depressive-like symptoms compared to mice maintained in light days with dark nights (LD). C57BL/6J mice received sham surgery or moderate T9 contusion SCI, then were placed permanently in LD or dLAN. dLAN after SCI did not worsen locomotor deficits; rather, SCI-dLAN mice showed slight improvement in open-field locomotion at the final timepoint. Although dLAN did not alter SCI-induced heat hyperalgesia, SCI-dLAN mice exhibited an increase in mechanical allodynia at 13 days post-SCI compared to SCI-LD mice. SCI-LD and SCI-dLAN mice had similar outcomes using sucrose preference (depressive-like) and open-field (anxiety-like) tests. At 21 dpo, SCI-dLAN mice had reduced preference for a novel juvenile compared to SCI-LD, implying that dLAN combined with SCI may worsen this mood-related behavior. Finally, lesion size was similar between SCI-LD and SCI-dLAN mice. Therefore, newly placing C57BL/6J mice in dLAN after SCI had modest effects on locomotor, pain-like, and mood-related behaviors. Future studies should consider whether clinically-relevant circadian disruptors, alone or in combination, could be ameliorated to enhance outcomes after SCI.

## Introduction

The Earth has rotated on its axis for billions of years, creating daily environmental rhythms of light and dark. These predictable daily light-dark cycles facilitated evolution of circadian rhythms in most organisms. Circadian rhythms are adaptive biological cycles that occur naturally over 24 hours to promote activity-rest patterns and general well-being (Blume et al., 2019). Entrainment of these cycles depends upon certain environmental cues, the strongest of which is light.

Rhythms in light exposure have been dampened and shifted in modern life, with reduced exposure to sunlight and increased artificial light at night (Lunn et al., 2017). Exposure to dim light at night (dLAN) is associated with health issues such as metabolic disruptions, oxidative stress, and immune system dysregulation (Fonken and Nelson, 2011, 2014). Circadian dysrhythmias are common in intensive care units (ICU) of hospitals, due to nighttime disruptions, inadequate exposure to daytime light, and constant exposure to low levels of light at night (Oldham et al., 2016). dLAN has detrimental effects on pathology and recovery in preclinical models. In mice, exposure to dLAN after stroke exacerbates negative outcomes and increased lesion size, which may have important implications for care of critically ill patients recovering in the ICU (Weil et al., 2020).

Spinal cord injury (SCI) affects approximately 300,000 people in the U.S. (Lasfargues et al., 1995). SCI elicits a hyperreactive response of immune cells, driving toxic neuroinflammation immediately following injury. Glial cells such as microglia, astrocytes, oligodendrocytes, oligodendrocyte progenitors, and blood-derived immune cells, play distinct roles in the SCI-elicited immune response. Although there are some positive effects of inflammation following SCI, extensive infiltration and overactivation of immune cells are major drivers of secondary damage after SCI. Cascading inflammation causes extensive cell death, deficits in locomotor activity, and chronic neuropathic pain (Gaudet and Fonken, 2018). In addition, individuals with SCI are predisposed to experiencing mood disorders like anxiety and depression (Lim et al.,2017). These behaviors may also be driven by inflammation (van West and Maes, 1999; Suarez et al., 2003). Given that the acute post-SCI inflammatory response likely exacerbates damage, strategies that quench early inflammatory reactions could benefit neuroprotection, recovery, and mood after SCI.

One potential neuroprotective pathway involves the link between circadian rhythms and inflammation. All known immune cells, including microglia and macrophages, contain intrinsic biological clocks, driving the regulation of immune responses (Keller et al., 2009; Nakazato et al., 2011). Indeed, “time-of-day" regulates microglia inflammatory reactivity; e.g., in rats, injection of lipopolysaccharide during the inactive phase induces more robust neuroinflammation compared injection during the active phase (Fonken et al., 2015). Circadian disruption also has detrimental effects in inflammatory diseases such as cancer, heart disease, and metabolic syndrome (Vyas et al., 2012; D’Ettorre et al., 2019). However, less is known about how circadian disruption affects neuroprotection and recovery after SCI.

Previous work explored the effects of dLAN on protein and gene expression to reveal disruption in core clock components (Fonken et al., 2013a). Swiss Webster mice exposed to ∼5 lux dLAN showed attenuated rhythmic expression of Bmal1, Per1, Per2, Cry1, and Cry2 mRNA relative to mice held on LD cycles. Additionally, dLAN reduced protein expression of both PER1 and PER2 in the SCN, revealing the effects of dLAN on gene- and tissue-specific changes in expression levels for various circadian clock genes. Exposure to dLAN increased microglial cytokine expression and sickness behavior following LPS administration (Fonken et al., 2013b), implying a possible effect of dLAN on inflammatory responses similar to that of SCI. Recent studies using C57BL/6J mice established that 5 lux dLAN can disrupt sleep patterns in young and aged mice(Panagiotou and Deboer, 2020), but had minimal behavior effect in a C57BL/6J derivative strain, B6.129S6-Per2tm1Jt/J (Cleary-Gaffney and Coogan, 2018). Further, C57BL/6J mice exposed to 15 lux dLAN during development had increased hippocampal inflammatory gene expression accompanied by anxiety and depressive-like symptoms in adulthood(Chen et al., 2021a). These studies suggest that this strain—commonly used for transgenic manipulation—is susceptible to circadian disruption via dLAN but may require higher levels of light exposure than other strains.

Here, we predicted that exposure to dLAN after SCI may amplify SCI-elicited immune responses within the injured spinal cord, thereby exacerbating secondary damage to the lesion site and worsening recovery after SCI. To test this hypothesis, mice were divided into SCI and Sham groups. SCI mice were subjected to T9 laminectomy and moderate (65 kDyn) contusion SCI; sham animals received T9 laminectomy only. Sham and SCI mice were subsequently housed under typical light-dark conditions (LD) or newly housed in dLAN for the remainder of the study. Animals from all four experimental groups were examined over a period of 35 days post-operative (dpo). To reveal the extent that aberrant light at night shifted biology and behavior after SCI, we assessed the effects of dLAN on locomotor recovery, neuropathic pain-like behaviors, anxiety- and depressive- like behaviors, and neuroprotection.

## Materials and Methods

### Animals

All housing, surgery, and postoperative care adhered to guidelines set by The University of Texas at Austin Institutional Animal Care and Use Committee. Adult middle-aged female and male C57BL/6J mice (∼30 weeks old; Jackson Laboratory, Stock 000664) were housed in sex-matched pairs. Food and filtered tap water were provided *ad libitum*.

### Housing and lighting conditions

Mice were maintained for 2+ weeks prior to surgery in cages in a lightproof Sun Calc Lighting Control System housing rack, which contains 14 different compartments with 6 cages per compartment. Each compartment is sealed and ventilated and has independently customizable light cycles. Thus, mice in different light treatment groups are maintained in otherwise identical conditions apart from a single variable – the light cycle. Sham and SCI mice in the two lighting groups were maintained in separate compartments, so that any possible odors or stressors differing between sham/SCI mice did not affect the other group. All mice were maintained on a 12 h bright light -12 h dark (LD) daily cycle prior to surgery. Sham-SCI and LD-dLAN groups of mice were divided based on pre-surgery mechanical and heat thresholds, such that the mean baseline thresholds between groups were as similar as possible. Immediately after surgery, mice were placed into two groups: (1) a typical 12:12 LD cycle; or (2) dLAN. dLAN consisted of 150 lux for 12 h during the daytime (same as bright light in the LD group) followed by 15 min of total darkness, then 15 lux light for 11.75 h during the nighttime (Chen et al., 2021a). dLAN was produced using LED light strips (Philips Hue). dLAN mice were placed in dLAN immediately after surgery and maintained in that lighting condition for the duration of the study (**Fig. 1a**).

**Figure 1.**
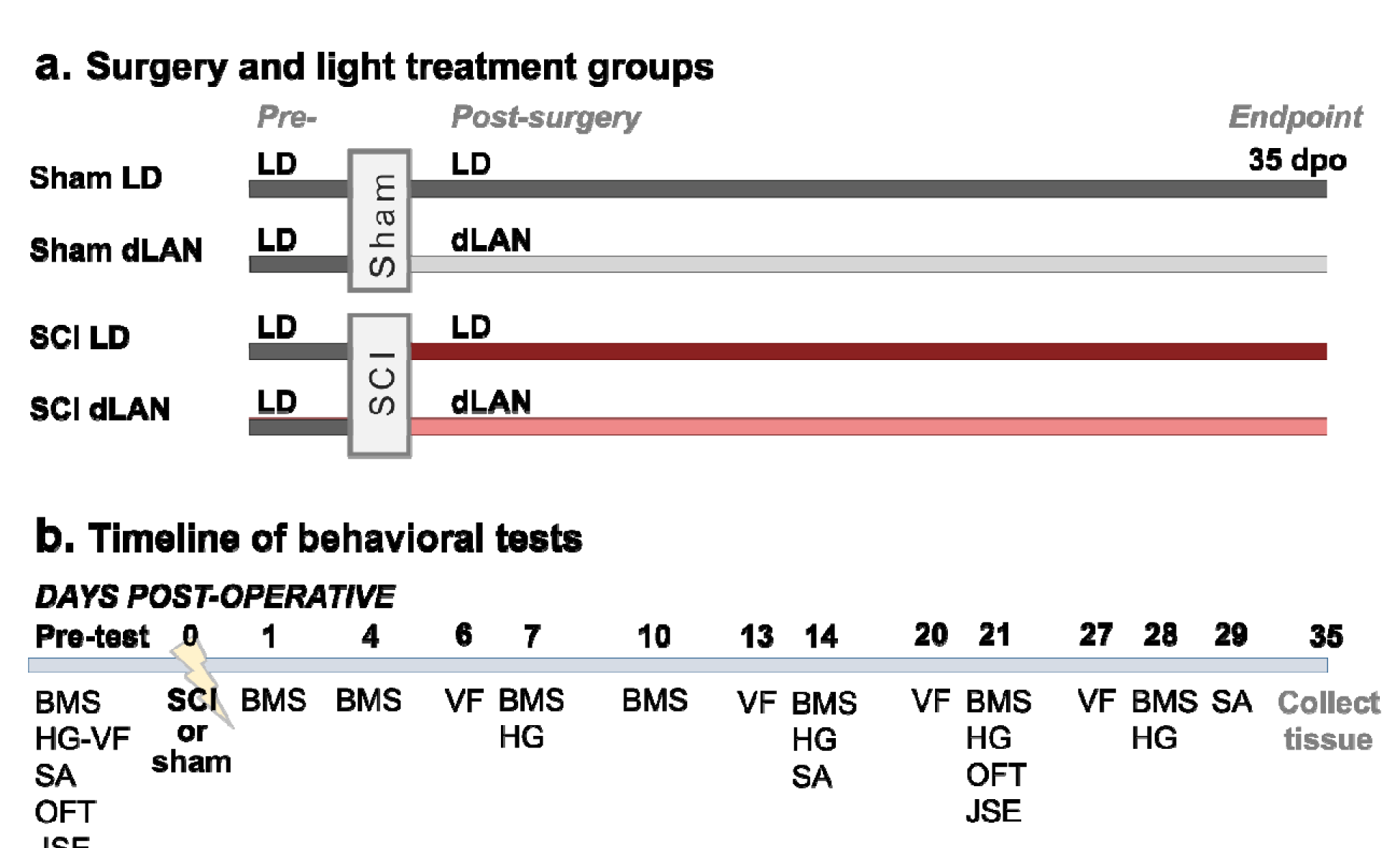
Experimental groups and timeline of behavioral tests. **a.** Surgery and treatment groups used in the study. Prior to surgery, all mice (females and males) were housed in light days and dark nights (LD: 12 h light:12 h dark). Mice received sham surgery or moderate 60 kDyn T9 contusion spinal cord injury (SCI). Immediately after surgery, mice were replaced in LD conditions or were newly housed in dim light-at-night (dLAN: light days with dim light-at-night). Mice were housed in these post-surgery lighting conditions for the remainder of the study. **b.** Schedule of testing for behavioral studies. Locomotor function was assessed repeatedly using the Basso Mouse Scale (BMS). Neuropathic pain-related behaviors were assessed weekly starting at 6-7 dpo using SUDO von Frey test (VF: mechanical allodynia) and Hargreaves test (HG: heat hyperalgesia). Anxiety-like symptoms were tested using the open-field test (OFT) and juvenile social exploration (JSE). Depressive-like symptoms were assessed using sucrose anhedonia (SA) and JSE. All outcomes were assessed prior to and after surgery; see Methods for more details.

The dLAN protocol employed in this study uses a higher intensity light (∼15 lux vs. ∼5 lux) than what has typically been used in previous mouse dLAN studies (Fonken et al., 2009, 2013a, 2013b, 2019; Weil et al., 2020). This decision was influenced by more recent studies suggesting that some C57BL/6-derived strains are less sensitive, in terms of circadian behaviors, to 5 lux dLAN, but that exposure to 15 lux dLAN during development can induce changes in behavioral and neuroimmune outcomes (Cleary-Gaffney and Coogan, 2018; Chen et al., 2021a). Further, 15 lux is also more in line with contemporary estimates of light pollution (Gaston et al., 2013; Walker et al., 2020) as well as observations from hospital settings (Engwall et al., 2015; Durrington et al., 2017; Nelson and DeVries, 2017).

### Surgery, mouse numbers, and animal care

All animals received buprenorphine hydrochloride (0.075 mg/kg; MWI Animal Health, Cat. 060969) analgesic immediately prior to surgery (to reduce acute post-surgery discomfort). Additional post-surgery analgesic was withheld to limit potential confounding effects on pain-related behavioral tests. T9 laminectomy was completed on both sham and SCI mice. Next, mice in SCI groups received contusion SCI using the Infinite Horizon impactor (Precision Systems and Instrumentation) (severity and forces defined below) (Scheff et al., 2003; Gaudet et al., 2016, 2021). Surgeries occurred between Zeitgeber time (ZT) 2-10 (lights on at ZT0 and off at ZT12). Incisions were closed using sutures and wound clips. Post-operative care comprised daily subcutaneous injections of Ringer’s solution (2, 2, 1, 1, 1 mL on the first 5 days post-operative (dpo), respectively; for both sham and SCI mice) to prevent dehydration, and manual voiding of bladders twice per day (shams handled similarly).

Mice received sham surgery (sham-LD: n=6 (4 females and 2 males); sham-dLAN: n=7 (3 females and 4 males)) or 65 kDyn T9 contusion SCI (SCI-LD: n=10 (5 females and 5 males), force: 67.5 ± 0.9 kDyn; displacement: 532 ± 17 μm) (SCI-dLAN: n=8 (6 females and 2 males), force: 68.3 ± 0.9 kDyn; displacement: 524 ± 12 μm). Mice were prospectively excluded from analysis if the Infinite Horizon force/displacement curves suggested a slip or bone hit; if SCI displacement was 100 μm higher or lower than the mean displacement; if mice died during surgery or prior to the experimental endpoint due to ill health/related euthanasia; or if the BMS score was unusually high at 1 dpo (BMS of 3 or greater) (Lee et al., 2023b). 5 male SCI mice were excluded (2 died during surgery, 1 died during bladder care, 1 due to bone hit, 1 due to high 1 dpo BMS score), 2 male sham mice were excluded (died during surgery), and 1 female sham mouse was excluded (died during surgery) (these mice are excluded from the final numbers presented above).

All animals received post-operative care and were replaced in LD conditions or newly placed in dLAN lighting conditions, as described above. Body mass was measured prior to and after surgery. SCI induced loss of body mass, which was not significantly affected by dLAN; however, SCI-dLAN mice did tend to have higher post-SCI body mass than SCI-LD mice (**Fig. 2**).

**Figure 2.**
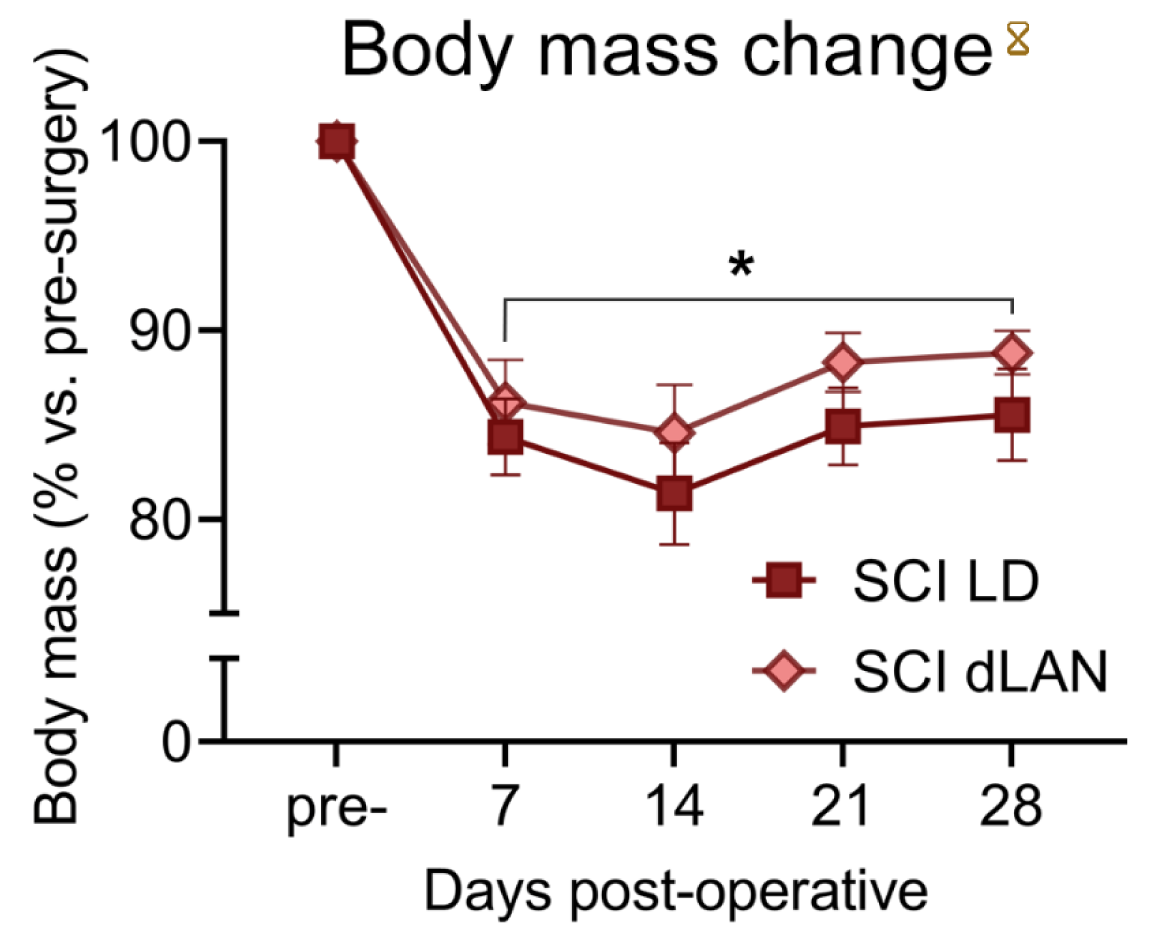
Body mass change in mice after SCI and placement in typical light-dark cycles (LD) or light days with dim light-at-night (dLAN). SCI reduced body mass in female and male mice. Lighting condition had no significant effect on body mass changes after SCI (*p* > 0.05). SCI-LD: n=10; SCI-dLAN: n=8. Plots show mean ± SEM. Hourglass symbol indicates *p* < 0.001 across timepoints (dpo main effect); * indicates *p* < 0.001 between pre-surgery and post-SCI body mass at all timepoints for both light treatments; two-way RM ANOVA with Bonferroni *post-hoc* test.

### Sex as a biological variable

Importantly, our study uses data from both male and female mice. Unfortunately, due to premature death or prospective exclusion too few males remained to draw meaningful conclusions regarding potential sex differences. For instance, analysis of locomotor recovery reveals little difference when we remove males entirely from the dataset (male & female: p = 0.50, F1,16 = 0.48; female only p = 0.54 F1,9 = 0.40; two-way RM ANOVA, light main effect) (see below). However, due to the unbalanced nature of the experimental groups, some differences observed in our study may result from underlying sex-differences.

### Locomotor testing

Behavioral tests and timing are summarized in **Fig. 1b**. Hindlimb motor function was measured using the Basso Mouse Scale (BMS) scores and subscores for locomotor recovery (Basso et al., 2006) prior to surgery and at 1, 4, 7, 10, 14, 21, and 28 dpo. Hindlimb recovery was scored on the BMS scale of 0 (complete paralysis) to 9 (normal motor function) by two observers blind to treatment conditions. BMS subscores range from 0 to 11. Subscore points are only counted once mice begin plantar stepping frequently or consistently; therefore, mice plantar stepping occasionally or not at all received a BMS subscore of zero.

### Neuropathic pain-related behavioral testing

Testing for neuropathic pain-associated behaviors was performed as previously described (Gaudet et al., 2017, 2021; Lee et al., 2023b). Mice were acclimated to the von Frey and Hargreaves apparatus for two 40-60 min sessions, followed by two pre-surgery tests (average of the two pre-surgery scores are presented here).

After surgery, sensory testing occurred weekly. We averaged values for the left and right hindlimb to calculate a single score for each animal at each timepoint. Testing occurred within 2 hours between ZT2-6. To control for potential confounds of having both sexes tested simultaneously, female and male mice were tested in separate sessions. Both sham and SCI mice from a single sex were tested in the same session; female and male cohorts were completed within 1-2 days of one another to minimize variability between groups and to enable meaningful comparisons. The order of mice in testing chambers was randomized to intersperse and remain blind to the treatment group.

*Von Frey test for mechanical allodynia:* Mechanical sensory thresholds were assessed using the simplified up-down (SUDO) method (Bonin et al., 2014) of von Frey testing to minimize stress and time outside of the home cage. Mice were placed in plastic compartments suspended above a wire mesh (Ugo Basile grid platform, Stoelting 57816) and acclimated for 40-60 min before testing. During testing, von Frey filaments (Touch Test Sensory Evaluator, Stoelting 58011) were applied to the center of the hindpaw plantar surface until the filament buckled and pressure was maintained for a maximum of 3 s. A positive response was established if the mouse withdrew their hindpaw or flinched in response to the filament pressure. To minimize mouse stress and ensure accurate scoring, von Frey testing occurred the day prior to Hargreaves testing.

*Hargreaves test for heat hyperalgesia:* Heat sensory thresholds (thermal hyperalgesia) were determined using the Hargreaves test (Ugo Basile Thermal Plantar test, Stoelting 55370) (Hargreaves et al., 1988; Cheah et al., 2017). Mice were maintained in a plastic box on top of a glass surface. An infrared heat source (intensity of 25) was applied to the center of the mouse’s hindpaw, and the latency to respond (lifting paw or flinching) was automatically recorded.

Maximum response latency was 30 s; the heat source automatically shut off at this time. Testing on each mouse’s right and left hindpaw was alternated (three tests per timepoint for a total of 6 tests), and each mouse had at least 5 min recovery between each test to minimize sensitivity.

### Anxiety- and depressive-related behavioral testing

*Sucrose preference test:* Sucrose anhedonia was assessed as described previously (Fonken et al., 2009; Chen et al., 2021b). Mice were first acclimated to novel 15 mL drinking tubes for two nights with a choice of water versus sucrose (sucrose bottle side was randomized initially then alternated upon subsequent trials) to acclimate them to the test. Mice were then presented a third time with the sucrose and water bottles prior to surgery for 14 h at the onset of the light phase. After surgery at 14 and 29 dpo, mice were again presented with the two-bottle choice overnight (14 h) beginning at the start of the active phase. Bottles were weighed prior to and after placement in the cages for every trial. Most mice were housed in sex-matched pairs, so the n for each sucrose preference result is the cage (i.e., results from two mice combined).

*Open field test:* Mice were tested on the open field test (Fonken et al., 2009; Chen et al., 2021a) prior to surgery and at 21 dpo. Mice were acclimated to the behavior room for 40 min. Open field tests were conducted during the light phase between Zeitgeber time (ZT) 4−9. Mice were placed in a random corner of an opaque plexiglass box (55cm × 55cm x 40 cm). Locomotor activity was recorded with EthoVision XT 14 (Noldus, Leesburg, VA) for 10 min. Distance traveled and the time in the center (center 30% of the whole arena) were automatically analyzed by the software over a 5 min period. The test box was cleaned with 70% ethanol after each trial. One female mouse from the SCI-LD group was excluded from open field analyses due to abnormally high central preference at 21 dpo (outlier according to Grubb’s test, *p* < 0.05).

*Juvenile Social Exploration:* Mice were tested on the juvenile social exploration (JSE) test (Chen et al., 2021a, 2021b) prior to surgery and at 21 dpo. Mice were acclimated to the behavior room for 40 min. Mice explored an open field apparatus for 10 min and the exploratory behaviors were scored automatically by EthoVision software, as described above. Immediately after the open field test, a same-sex novel juvenile stimulus mouse (∼4 weeks old) was introduced in the arena for a 5 min JSE test. An experienced scorer who was blinded to the experimental groups analyzed social exploration (sniffing, grooming, chasing, and fighting) initiated by the experimental mouse using EthoVision software.

### Immunohistochemistry and lesion analysis

*Tissue collection and cryosectioning:* at 35 dpo, mice were euthanized via an intraperitoneal injection of Euthasol (200-270 mg/kg; Virbac AH, Inc.) followed by transcardial perfusion with cold 1X Phosphate Buffered Saline (PBS; pH 7.4) and 4% paraformaldehyde in 1X PBS. Spinal cords (5 mm segments including the injury epicenter and 2.5 mm rostral and caudal) were dissected and immediately placed in cold 4% paraformaldehyde overnight before being transferred to 30% sucrose the next day. Spinal cords were blocked in optimal cutting temperature (OCT; Fisherbrand) compound and flash frozen on dry ice. A cryostat was used to obtain 10 µm serial cross sections which were collected onto Superfrost Plus slides (Fisherbrand) and stored at -20°C for later use. LD and dLAN mice were randomized so that tissue collection, handling, and subsequent analysis were all performed in a blind manner.

*Immunohistochemistry* was performed as previously described (Gaudet et al., 2016). Slides were first brought to room temperature and then a hydrophobic barrier pen was used to encircle the sections. Slides were washed in 1X PBS for 10 min and then blocked with 10% normal donkey serum (NDS) in 1X PBS with 0.2% Triton X-100 (PBTx) for 1 hr at room temperature in a humidified chamber. The following primary antibodies were diluted in 10% NDS in PBTx and applied overnight: Rabbit anti-Iba1 (1:1,000; Wako; RRID: AB_839504) and Rat anti-GFAP (1:500; Invitrogen; RRID: AB_2532994). The next day, slides were washed in PBS three times for 5 min at room temperature followed by incubation at room temperature for 2 hrs with the following secondary antibodies and DAPI diluted in PBTx: Donkey anti-Rabbit IgG Alexa Fluor 568 (1:500; Thermo Fisher; RRID: AB_2534017) and Donkey anti-Rat IgG Alexa Fluor 488 (1:500; Thermo Fisher; RRID: AB_2535794). Finally, slides were washed three times in PBS for 5 min at room temperature before being mounted in Immu-Mount (epredia), covered with a coverslip, and sealed with nail polish. Immunohistochemistry to analyze white matter sparing followed a similar procedure but using a Goat anti-NF-H antibody (1:3000; Thermo Fisher PA5- 143591) along with Donkey anti-Goat IgG Alexa Fluor 568 (1:500; Thermo Fisher; RRID:

AB_2762828) to identify Neurofilament heavy chain immunoreactivity. After secondary antibody washes, 1x FluoroMyelin Green (Thermo Fisher; F34651) was applied for 30 minutes followed by 3 x 5min PBS washes then mounting as before.

*Imaging and lesion analysis*: spinal cord sections were imaged on a Nikon Ni-E widefield epifluorescence microscope using the “*acquire large image*” application in Nikon’s *Elements* imaging software (AR 5.21.02). Nine overlapping (20%) 10x fields were acquired per cross section and digitally stitched together to form a single large image. Epicenter sections, defined as the section with the largest lesion by area, and 10 surrounding sections (5 caudal and 5 rostral) spaced 200 µm apart were acquired per mouse. Images were analyzed in *ImageJ* (v2.1.0 *Fiji* distribution) (Schindelin et al., 2012; Schneider et al., 2012) to determine lesion area and total area based on GFAP staining. White matter area was determined based on NF-H immunoreactivity and FluoroMyelin presence, excluding NF-H+ grey matter. Data was decoded, organized, and matched to specific mice using the following *R* (v4.2.2) (R Core Team, 2022) packages via *RStudio* (2023.03.1 build 446) (RStudio Team, 2020): *tidyverse* (v2.0.0) (Wickham et al., 2019; Wickham, 2023), *splitstackshape* (1.4.8) (Mahto, 2019), and *readxl* (v1.4.2) (Wickham and Bryan, 2023).

### Statistics

Data were analyzed using two-way or three-way repeated measures (RM) ANOVA and Holm-Sidak *post-hoc* tests where appropriate using *SigmaPlot* (v15.0). Tests used are stated in the text for each dataset. Data were graphed using *GraphPad Prism* (v9.5.1).

Statistical analysis was performed using SigmaPlot 15.0, and results with *p* < 0.05 were defined as significant. All data are plotted as mean ± SEM.

Study data were deposited at the Open Data Commons for Spinal Cord Injury (ODC-SCI; RRID:SCR_016673). Accession: 956. The ODC-SCI is a secure, cloud-based repository platform designed to share research data (Torres-Espín et al., 2022; Aldrich et al., 2023)

## Results

### Effects of dim light-at-night on locomotor recovery after SCI

We sought to establish whether dLAN worsened outcomes after SCI in adult C57BL/6J mice. Prior to surgery, all mice were maintained in LD conditions. Mice then received surgery (laminectomy sham or moderate T9 contusion SCI); next, mice were immediately housed in LD (control conditions) or in dLAN for the rest of the study. Locomotor function was assessed using the BMS and BMS subscore (Basso et al., 2006). After SCI, both SCI-LD and SCI-dLAN mice displayed expected locomotor deficits, which recovered over 28 dpo (**Fig. 3**) (three-way ANOVA, surgery x dpo interaction: *F*_7,216_ = 79.18, *p* < 0.001). There were no significant differences in locomotor recovery between LD and dLAN mice (two-way RM ANOVA, light x dpo interaction: *F_7,112_* = 1.87, *p* = 0.08), except at the final timepoint (28 dpo) when dLAN mice had modestly enhanced locomotor function according to the BMS score and BMS subscore (Holm-Sidak *post-hoc* test, LD-SCI vs. dLAN-SCI: *p* < 0.05). Therefore, dLAN did not exacerbate SCI-elicited locomotor deficits in C57BL/6J mice compared to LD as predicted; rather, dLAN may have slightly improved locomotor recovery after SCI.

**Figure 3.**
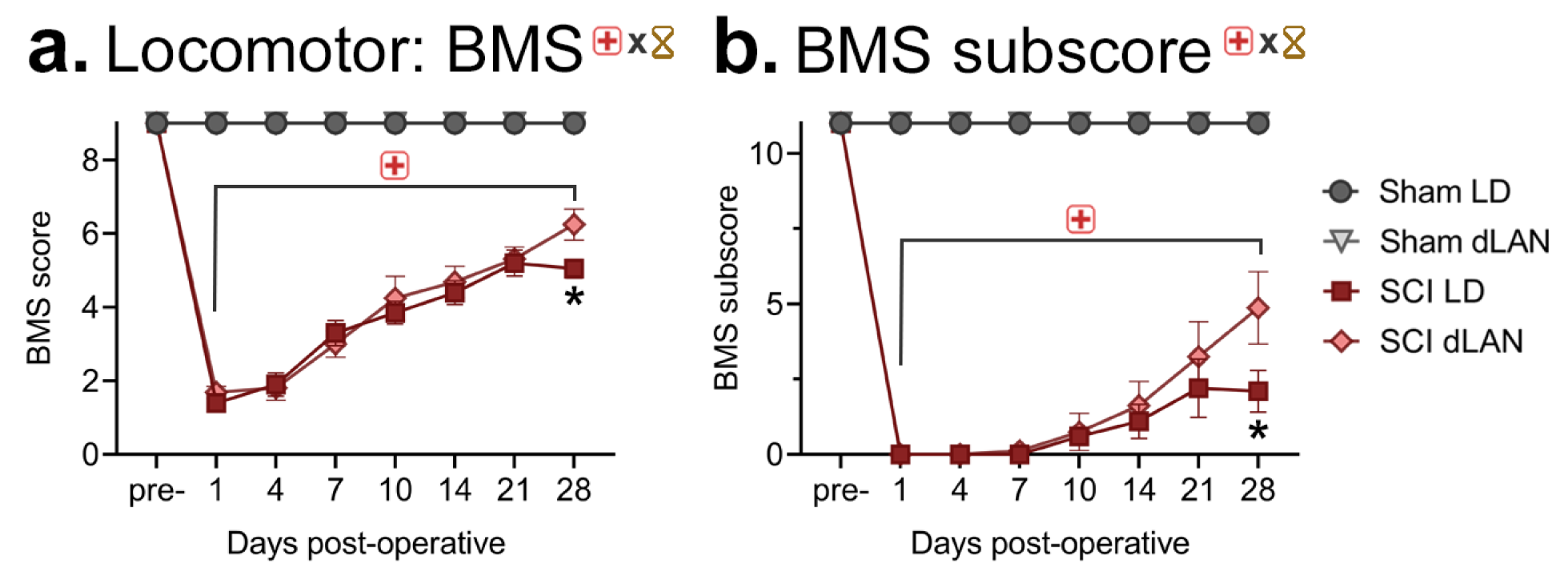
Effects of dLAN on locomotor recovery after SCI as measured by the Basso Mouse Scale (BMS) score and subscore. **A.** BMS scores showing SCI-elicited hindlimb locomotor deficits, which recovered over time. **B.** BMS subscores enabled assessing more detailed recovery of walking patterns after SCI. SCI mice exposed to dLAN had higher BMS scores and subscores at 28 dpo compared to SCI-LD mice. Sham-LD: n=6; Sham-dLAN: n=7; SCI-LD: n=10; SCI-dLAN: n=8. Plots show mean ± SEM. Significant (*p* < 0.05) surgery (red cross) x dpo (hourglass) interactions were observed via a three-way ANOVA. Timepoints with significantly different SCI vs. Sham results—as determined by Holm-Sidak post-hoc testing— are indicated with a red cross and brackets, while significant SCI-LD vs SCI-dLAN differences are indicated by an asterisk.

### Effects of dim light-at-night on SCI-elicited pain-related behaviors

In addition to effects on locomotor recovery, we also examined the effects of dLAN on SCI-elicited pain-like behaviors using Von Frey and Hargreaves testing weekly after injury (Hargreaves et al., 1988; Bonin et al., 2014; Cheah et al., 2017). Hargreaves testing revealed that mice with SCI experienced heat hypersensitivity compared with sham mice (three-way ANOVA, surgery x dpo interaction: *F_4,133_* = 17.29, *p* < 0.001); however, exposure to dLAN had no significant effect on sensitivity to thermal stimuli when compared with mice in LD (control) conditions in sham or SCI mice (light x surgery x dpo interaction: *F_4,133_*= 0.808, *p* = 0.522) (**Fig. 4a, b**). Von Frey assessment for mechanical sensitivity similarly showed hypersensitivity in mice subjected to SCI at all post-operative timepoints (three-way ANOVA, surgery x dpo interaction: *F_4,135_* = 4.629, *p* = 0.002). Interestingly, SCI mice exposed to dLAN had exacerbated mechanical hypersensitivity at 13 dpo, when compared with those animals kept in LD conditions (Holm-Sidak *post-hoc* test, LD-SCI vs. dLAN-SCI: *p* < 0.05), although this effect was transient as the groups are indistinguishable at subsequent timepoints (**Fig. 4c,d**). Together, these data suggest that *de novo* exposure to dLAN after SCI has no significant effect on heat sensory thresholds, and that dLAN modestly worsens SCI-induced mechanical hypersensitivity.

**Figure 4.**
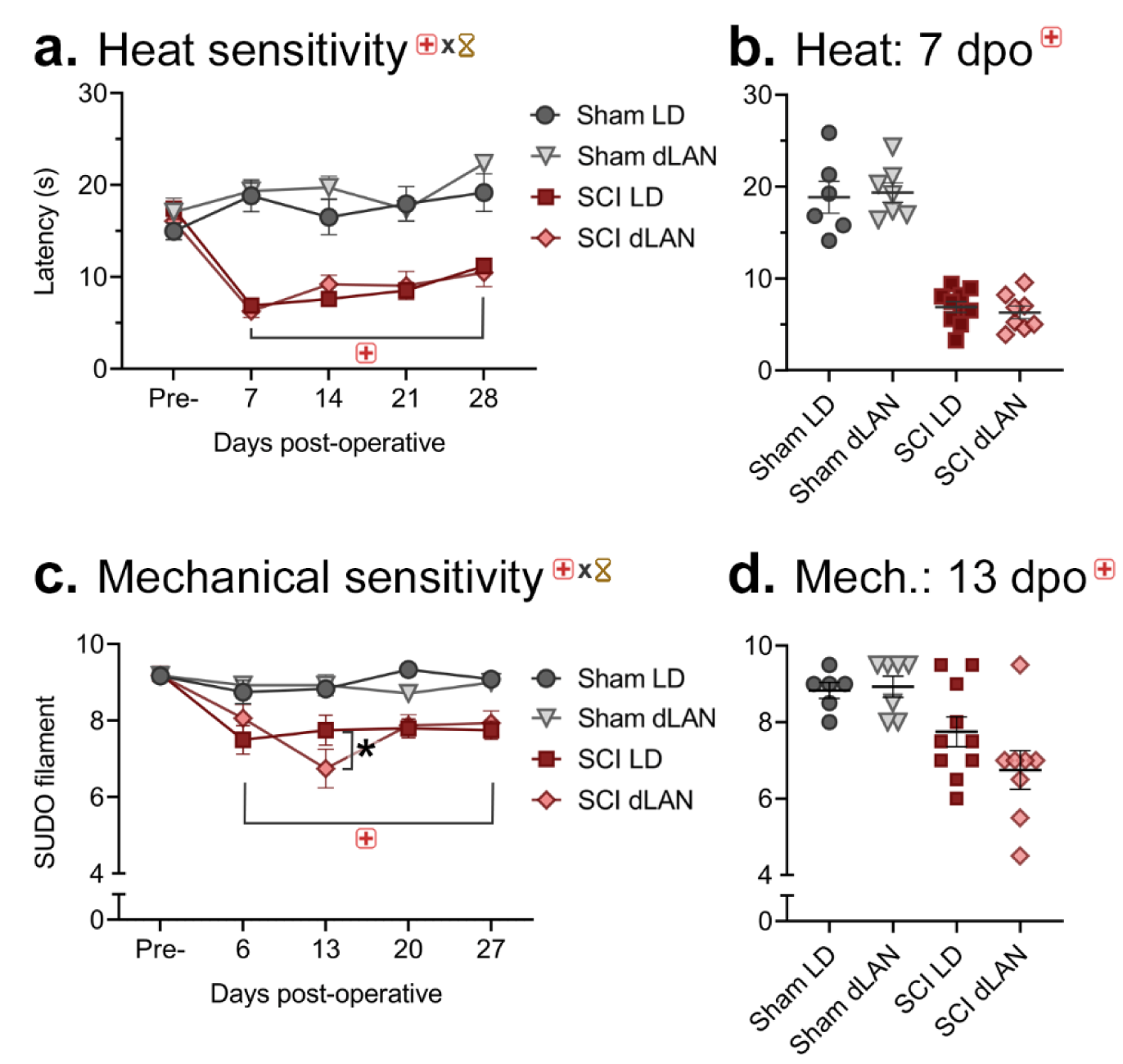
Effects of dLAN on contusion SCI-mediated neuropathic pain symptoms. **A.** SCI induced hypersensitivity to heat from 7-28 dpo as assessed by the Hargreaves test. There was no significant difference in heat sensitivity between LD or dLAN groups. **B.** Plot of individual withdrawal latencies to heat at 7 dpo to illustrate post-surgery difference of SCI vs Sham animals. **C.** SCI elicited hypersensitivity to mechanical stimulation as assessed by the von Frey test. SCI-dLAN mice showed a transient increase in mechanical sensitivity at 13 dpo compared to SCI-LD mice. **D.** Plot of individual SUDO filament thresholds at 13 dpo. Sham-LD: n=6; Sham- dLAN: n=7; SCI-LD: n=10; SCI-dLAN: n=8. Plots show mean ± SEM. Significant (*p* < 0.05) surgery (red cross) x dpo (hourglass) interactions were observed via a three-way ANOVA. Timepoints with significantly different SCI vs. Sham results—as determined by Holm-Sidak post-hoc testing—are indicated with a red cross, while significant SCI-LD vs SCI-dLAN differences are indicated by an asterisk.

### Effects of dim light at night on anxiety- and depressive-like behaviors

To explore the effects of dLAN on anxiety- and depressive-like behaviors, three related behavioral assays were conducted on mice housed after surgery in typical LD conditions or dLAN.

We assessed depressive-like behavior using the sucrose preference test: mice in this test typically show strong preference for sweet sucrose over water; reduced preference after surgery or treatments are interpreted as anhedonia (Christianson et al., 2008; Chen et al., 2021b).

Individual cages, consisting of sex-matched pairs, were assessed for sucrose anhedonia prior to surgery, and again at both 14 and 29 dpo. Our data revealed no significant effect of either SCI or dLAN on sucrose preference at 14 or 29 dpo (three-way ANOVA, light x surgery x dpo interaction: *F*_2,42_ = 0.00755, *p* = 0.992) (**Fig 5a-c**).

**Figure 5.**
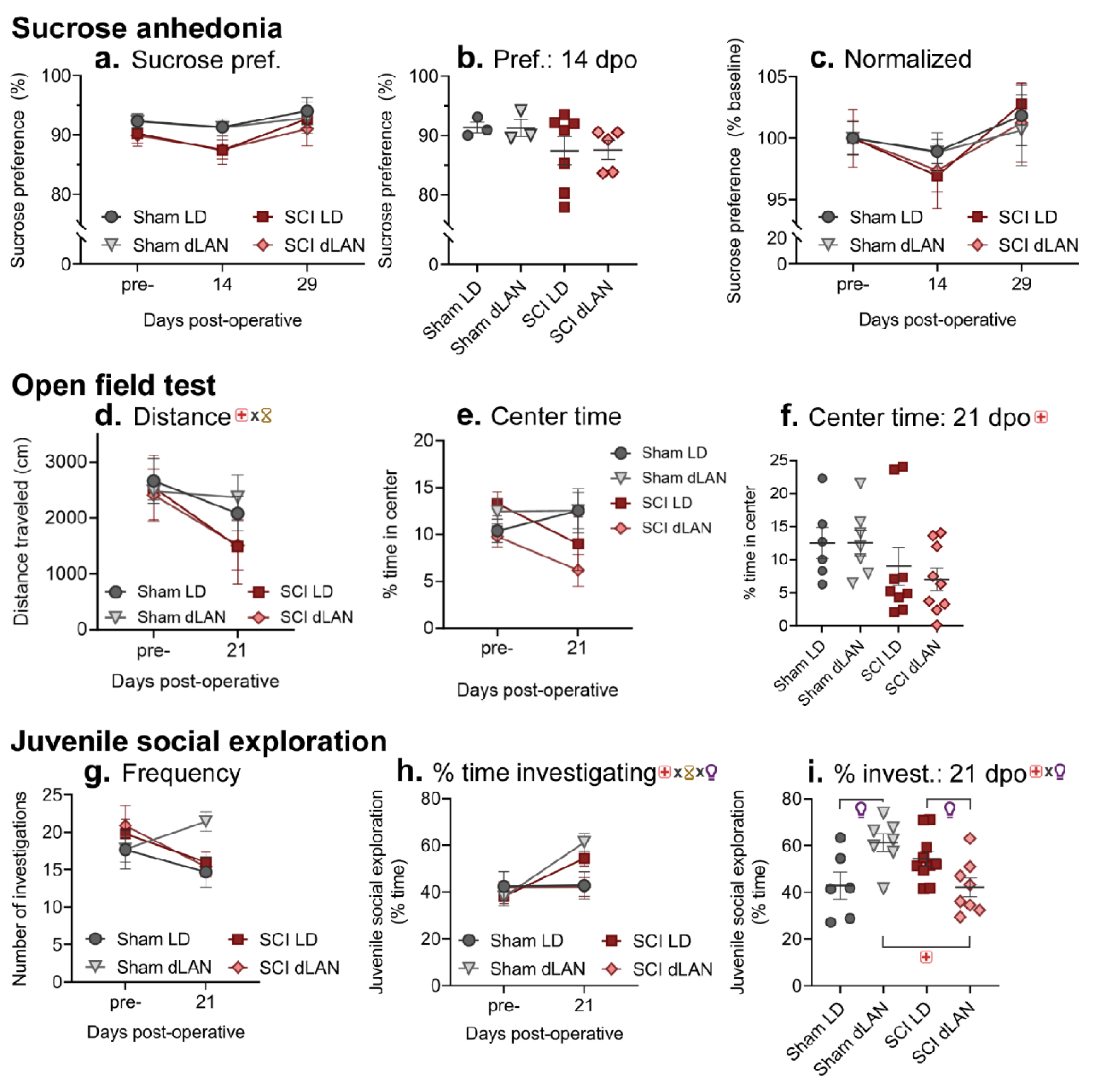
Effects of SCI and dLAN on depressive- and anxiety-like behaviors. **A-c**. Sucrose anhedonia was used to assess depressive-like behavior prior to surgery and at 14 and 29 dpo. Neither SCI nor dLAN significantly altered sucrose preference. **D-f.** Open field test: Mice were placed in an open field for 5 minutes to assess locomotion and anxiety-like behavior. **D.** At 21 dpo, SCI mice traveled less distance than sham mice. **E-f.** Mice with SCI displayed reduced central tendency at 21 dpo compared to sham mice. **G-i.** Juvenile social exploration: Mice in an open field were introduced to a novel juvenile; frequency and time of investigation were assessed as measures of anxiety- and depressive-like behavior. **H.** Mice showed similar behavior at baseline, whereas social exploration was significantly altered by lighting and SCI. **i.** At 21 dpo, sham-dLAN mice had increased exploration compared to sham-LD mice and SCI- dLAN mice. In addition, SCI-LD mice had increased exploration compared to SCI-dLAN mice. Sham-LD: n=6; Sham-dLAN: n=7; SCI-LD: n=9-10; SCI-dLAN: n=9. Plots show mean ± SEM. Significant (*p* < 0.05) main or interaction (indicated by “x”) effects from two or three-way ANOVA tests are indicated by red cross (surgery), hourglass (dpo), and light bulb (light condition) symbols at the top of each panel while significant Holm-Sidak *post-hoc* test results are indicated by brackets where applicable.

We examined anxiety-like behavior using the open field test. Mice with increased anxiety-like behavior are expected to decrease time in the center of the open field (Fonken et al., 2009; Chen et al., 2021a). Prior to surgery, mice from all groups had similar distance traveled and percent time in center (**Fig. 5d,e**). Overall, little difference was observed in percent time in the center zone between pre-surgery and post-surgery tests (**Fig. 5e**), though SCI mice, regardless of light treatment, spent less time in the center compared to sham mice at 21 dpo (two-way ANOVA, surgery main effect: *F_1,26_* = 4.310, *p* < 0.048) (**Fig. 5f**). Although decreased center zone time is typically considered increased anxiety-like behavior, SCI mice also travel less distance (three-way ANOVA, surgery x dpo interaction: *F*_1,52_ = 11.114, *p* = 0.002) (**Fig.5d**). Therefore, further testing is needed to disentangle SCI-induced differences in center zone time observed at 21 dpo. No significant dLAN effect was observed in distance traveled (three-way ANOVA, light x surgery x dpo interaction: *F*_1,52_ = 0.411, *p* = 0.524) or center zone time at 21 dpo (two-way ANOVA, light main effect: *F*_1,26_= 0.346, *p* = 0.561), suggesting that dLAN does not alter post-SCI changes in open field test behavior.

The JSE test was also used to assess anxiety- and depressive-like behavior. Mice typically spend substantial time exploring a novel, non-threatening juvenile mouse; mice with anxiety- or depressive-like behaviors are expected to decrease exploration of the novel juvenile (Christianson et al., 2008). Pre-surgery, all mice showed similar behavior (**Fig. 5g,h**); however, at 21 dpo, both lighting and injury altered percent time investigating a novel juvenile (two-way ANOVA, surgery x light interaction: *F_1,27_*=13.685, *p*<0.001) (**Fig. 5i**). Sham-dLAN mice increased time investigating compared to Sham-LD mice (Holm-Sidak *post-hoc* test, *p* = 0.007). In contrast, SCI-dLAN mice had decreased time exploring the juvenile compared to both SCI-LD (*p* = 0.031) and sham-dLAN mice (*p* = 0.003). Therefore, dLAN after SCI decreased time investigating a novel juvenile, suggesting that dLAN may elicit anxiety- and/or depressive-like behaviors as assessed using the JSE test.

### Effects of dim light-at-night on lesion size and neuroprotection at 35 dpo

Finally, to assess the extent of post-SCI tissue damage in dLAN vs LD housed mice, we collected spinal cord segments from a subset of mice with SCI (9 SCI-LD and 7 SCI-dLAN mice) and used immunofluorescence to analyze lesion size and tissue sparing. We defined the lesion border based on anti-GFAP immunoreactivity of astrocytes (**Fig. 6a**) and measured the lesion area at the epicenter and at 10 adjacent evenly spaced positions (**Fig. 6b**). We observed no significant difference between LD and dLAN overall (two-way RM ANOVA, light effect: *F_1,14_* = 0.269, *p* = 0.612) or at any specific rostro-caudal position (light x position effect: *F_10,138_* = 0.625, *p* = 0.790). We also quantified the degree of neuroprotection under each lighting condition by calculating the percentage of spared tissue—i.e., we subtracted the lesion area from the total cross-sectional area and expressed this as a percentage (**Fig. 6c**). In support of our lesion area measurements, we again saw no significant difference between LD and dLAN overall (two-way RM ANOVA, light effect: *F_1,14_*= 1.025, *p* = 0.329) or at any specific position (light x position effect: *F_10,138_* = 0.556, *p* = 0.847). To compliment these findings, we identified white matter with a combination of FluoroMyelin Green and anti-Neurofilament H immunohistochemistry (**Supplemental Fig. 1a**) and quantified the spared white matter as a percentage of total cross-sectional area at the epicenter and six adjacent positions (**Supplemental Fig. 1c**). No difference significance was observed between LD and dLAN overall (two-way RM ANOVA, light effect: *F_1,14_* = 2.35, *p* = 0.148) or at any specific rostro-caudal position (light x position effect: *F_6,84_* = 0.97, *p* = 0.453). Together, these anatomical results corroborate the locomotor recovery findings presented above, suggesting that dLAN does not exacerbate post-SCI neurotoxicity or locomotor recovery in C57BL/6J mice.

**Figure 6.**
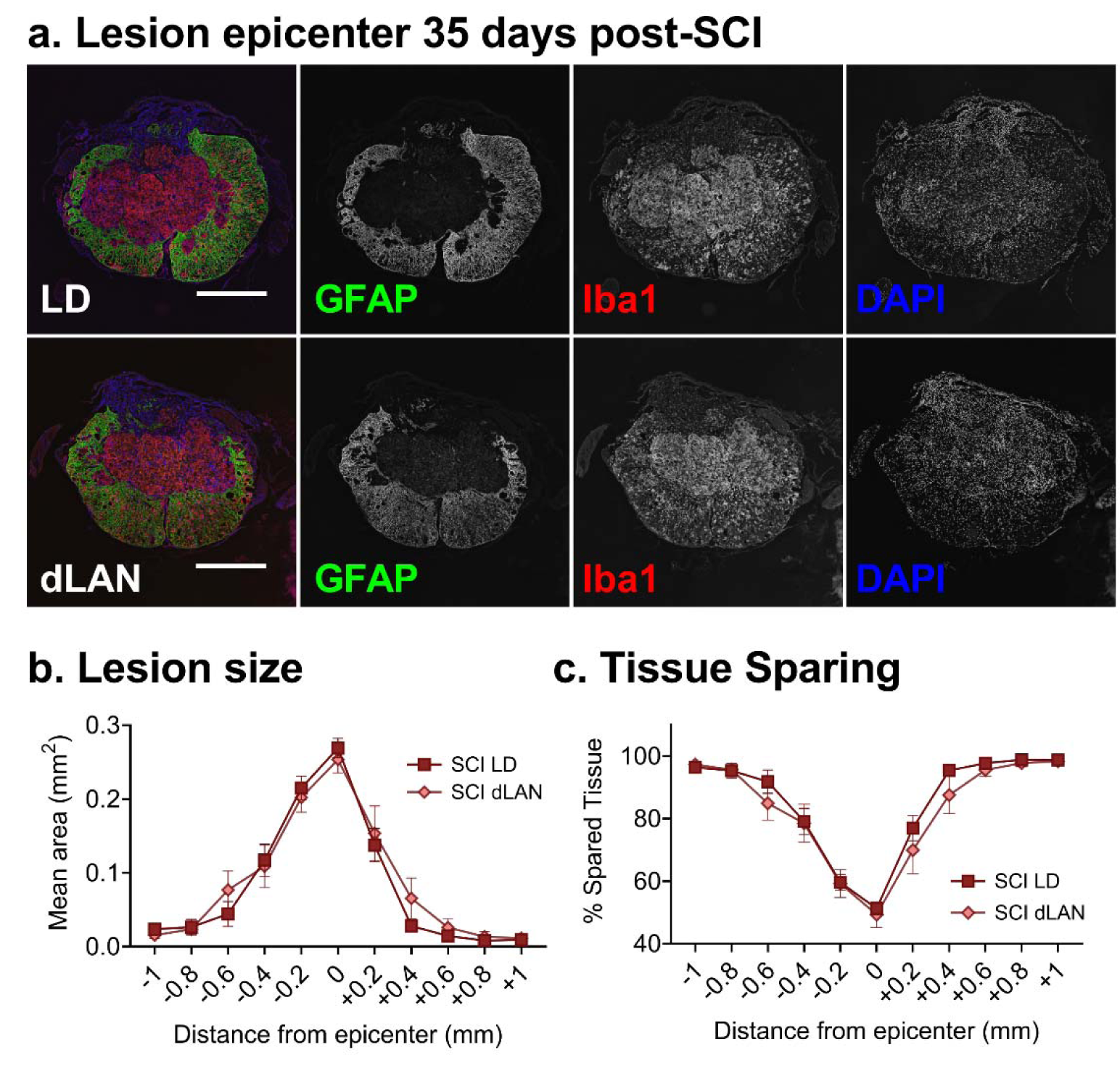
dLAN did not significantly alter post-SCI lesion size and tissue sparing in C57BL/6J mice. 10 µm spinal cord cryosections from 35 dpo mice were imaged at regular 0.2 mm intervals up to -1 mm rostral and +1 mm caudal of the lesion epicenter. **A.** Representative epicenter spinal cord cross-sections from SCI mice exposed to LD or dLAN. Sections were immunostained with anti-GFAP and anti-Iba1 to detect reactive astrocytes and microglia/macrophages respectively, along with DAPI to counterstain nuclei. Lesion area was determined by manually tracing the border based on anti-GFAP staining. **B.** dLAN after SCI had no significant effect on lesion size nor tissue sparing. Spared tissue was determined by subtracting the lesion area from the total area of the spinal cord section—this is expressed as the percentage of spared tissue relative to the total area. Lighting condition had no significant effect on either lesion area or tissue sparing (p = 0.61 and 0.33 respectively) based on two-way repeated measures ANOVA. SCI-LD: n=9; SCI-dLAN: n=7. Plots show mean ± SEM. Scale bars represent 500 µm.

## Discussion

Circadian disruption and suboptimal lighting is common in hospital settings (Tamburri et al., 2004; Gazendam et al., 2013; Oldham et al., 2016). Circadian disruption and dLAN also worsens outcomes in preclinical models of neurological disorders (Fonken et al., 2019; Weil et al., 2020; Hetman et al., 2022), yet few studies have addressed whether clock-related disruption exacerbates recovery after SCI (Slomnicki et al., 2020, 2021). To test this possibility, we subjected C57BL/6J mice raised in standard LD conditions (12 h light (150 lux); 12 h dark) to SCI or sham surgeries and then subsequently housed them in LD or dLAN (12 h 150 lux light (150 lux); 12 hr dim light (15 lux)) conditions (**Fig. 1**). We found that subjecting C57BL/6J mice to SCI then placing them in dLAN had some modest effects on locomotor and pain-related outcomes: SCI mice subjected to dLAN had slightly better locomotor function at 28 dpo. SCI- dLAN mice had worsened mechanical hypersensitivity at 13 dpo, compared to SCI-LD mice.

SCI-dLAN mouse behavior was similar to SCI-LD mice on the sucrose preference test (depressive-like behavior) and on the open-field test (anxiety-like behavior). Interestingly, SCI- dLAN mice had reduced juvenile social exploration compared to SCI-LD mice, which may indicate increased depressive-like behavior. Overall, our data suggest that dLAN after SCI in C57BL/6J mice slightly improves locomotor function and modestly worsens mechanical hypersensitivity and depressive-like behavior; implications for future preclinical studies and clinical translation are discussed below.

Our central hypothesis—that dLAN would exacerbate post-SCI symptoms and recovery outcomes—was prompted by a pair of studies that explored similar questions in mouse models of global cerebral ischemia and ischemic stroke (Fonken et al., 2019; Weil et al., 2020). In both studies, housing mice in dLAN post-operation resulted in increased neuronal cell death accompanied by the upregulation of pro-inflammatory factors (e.g. *Tnf*, *IL6*, and *IL1b*). These findings are consistent with a larger body of work demonstrating the relationship between circadian rhythms and the neuroinflammatory system (Fonken et al., 2015, 2016b, 2016a; Wang et al., 2020; Guzmán-Ruiz et al., 2023).

Given the overall importance of neuroinflammation in driving post-SCI secondary damage (Stirling and Yong, 2008; Karimi-Abdolrezaee and Billakanti, 2012; Gensel and Zhang, 2015; Ahuja et al., 2017; Gaudet and Fonken, 2018), it seemed likely that disruption of the circadian system via altering exogenous light conditions would inhibit post-SCI hindlimb locomotor recovery. BMS (Basso et al., 2006) was used to assess locomotor recovery at various timepoints from 1-28 dpo and very little overall difference was observed between dLAN and LD housed SCI mice (**Fig. 3**). If anything, we observed a small but statistically significant improvement in dLAN mice at 28 dpo, which is in contrast to dLAN results in other models of CNS injury (Fonken et al., 2019; Weil et al., 2020). Although—to our knowledge—no previous study has specifically tested effects of dLAN on SCI, a pair of recent studies has examined related circadian interventions (Slomnicki et al., 2020, 2021). Slomnicki *et al*. (2020) investigated post-SCI recovery and neuroprotection in mice lacking *Bmal1*—a core circadian clock component (Bunger et al., 2000; Bell-Pedersen et al., 2005)—and found a beneficial effect. In addition to uncovering a potential role for *Bmal1* in enhancing the neuroinflammatory response after SCI, these findings also illustrate the fact that disruptions to the circadian clock— depending on the specific experimental context—are not intrinsically harmful. In another study, the same group examined how time of injury effects post-SCI recovery and found little to no effect (Slomnicki et al., 2021), in contrast with previous work demonstrating that the neuroimmune system is more reactive during the light phase (Weil et al., 2009; Wang et al., 2020). Notably, this work used C57BL/6J mice, which may be important (as discussed below).

65-80% of individuals with SCI experience chronic neuropathic pain (Siddall et al., 1999) and pain-associated behaviors are similarly reported post-injury in various SCI models (Detloff et al., 2008; Putatunda et al., 2014; Gaudet et al., 2017, 2021). We used the Hargreaves test (Hargreaves et al., 1988; Cheah et al., 2017) and the SUDO von Frey approach (Bonin et al., 2014) to test the effects of dLAN on heat and mechanical hypersensitivity respectively (**Fig. 4**). As previously reported (Gaudet et al., 2017; Lee et al., 2023b), moderate T9 contusion SCI induces post-operative heat hyperalgesia and mechanical allodynia relative to sham surgery. We established that dLAN elicited a decrease in the SUDO threshold in SCI mice at 13 dpo— indicating that dLAN exacerbated SCI-induced mechanical allodynia at that timepoint (**Fig. 4c, d**). Like secondary damage at the lesion epicenter, SCI induced neuropathic pain is closely linked to heightened neuroinflammation. In the specific context of a thoracic injury model, remote activation of immune cells in regions caudal to the site of injury lead to neuronal damage and hindlimb hypersensitivity (Suter et al., 2007; Detloff et al., 2008; Grace et al., 2014; Watson et al., 2014). Our observation that post-SCI dLAN has little effect on neuropathic pain is consistent with our BMS locomotor recovery results (**Fig. 3**) which broadly suggest that exposure to dLAN after SCI may not meaningfully enhance the neuroimmune response in C57BL/6J mice.

Individuals with SCI commonly experience a higher prevalence of anxiety, depression, and various mood disorders compared to the general population (Kessler et al., 2012; Lim et al., 2017; Peterson et al., 2020). Likewise, post-SCI anxiety-like and depression-like symptoms occur in clinically-relevant rodent models (Luedtke et al., 2014; Farrell and Houle, 2019; Fukutoku et al., 2020; Lee et al., 2023a) with neuroinflammation as a key factor (Wu et al., 2014b, 2014a; Maldonado-Bouchard et al., 2016; do Espírito Santo et al., 2019; Brakel et al., 2021). Furthermore, dLAN and other circadian disruptions can induce changes in anxiety-like and depression-like behaviors independently of any surgical intervention (Bedrosian et al., 2013; Borniger et al., 2014; Cissé et al., 2016; Chen et al., 2021b, 2021a). In the JSE test, we found that dLAN decreased anxiety-like behavior in sham mice—i.e., social exploration increased (Christianson et al., 2008)—while the opposite effect was observed in SCI mice where dLAN-SCI mice spent less time on juvenile investigation compared to LD-SCI mice (**Fig. 5g-i**). Decreased anxiety-like behavior in sham mice exposed to dLAN vs. LD is consistent with some previous findings in rodents, depending on the test used (Bedrosian et al., 2011; Aubrecht et al., 2013; Hogan et al., 2015; Russart and Nelson, 2018). The SCI results are interesting in suggesting that dLAN interacts with SCI to enhance anxiety-like behavior. Previous studies— using various testing paradigms—have suggested that SCI alone amplifies anxiety-like behaviors (Fukutoku et al., 2020; Lee et al., 2023a) including a reduction in juvenile social interactions in rats (do Espírito Santo et al., 2019), whereas our results with JSE show that SCI mice housed in LD have similar anxiety-like behaviors as sham-LD mice. Further, mice with SCI exposed to dLAN display amplified anxiety-like behavior compared to SCI-LD mice (**Fig. 5h**).

Differences in anxiety-like behavior outcomes compared to previous studies likely relate to timing of tests (dpo); surgery model; and a potential learning effect from prior testing. Overall, results from the sucrose preference test and open field test showed little effect of dLAN on post-SCI depressive- or anxiety-like behaviors, while noting the following caveats. First, the sucrose preference test results are an average of sex-matched pairs (see Methods), which may not be as sensitive in distinguishing individual differences compared to the other behavioral assays.

Second, post-SCI JSE and OFT took place at 21 dpo, before the BMS locomotor difference was noted at 28 dpo (**Fig. 3**). dLAN-elicited effects may have been different or more pronounced if these tests had been conducted at a later timepoint.

Neuroinflammation drives secondary damage after SCI and the degree of post-SCI locomotor recovery is closely linked to the amount of spared white matter in and around the lesion epicenter (Gaudet and Fonken, 2018). We quantified the effect of post-surgery light treatment on lesion size and spared tissue by examining immunostained cryosections taken from the lesion epicenter and surrounding tissue of 35 dpo mice (**Fig. 6**). We observed no difference between light treatments, suggesting that dLAN does not significantly enhance the post-SCI neuroinflammatory response in C57BL/6J mice. Given the noted correlation between locomotor recovery and white matter sparing (Ma et al., 2001; Schucht et al., 2002; Basso et al., 2006), these results are consistent with our BMS results (**Fig. 3**) which show minimal effects of dLAN on post-SCI functional recovery in this strain. Similarly, SCI induced neuropathic pain is linked to heightened neuroinflammation. Remote activation of immune cells in regions caudal to thoracic SCI lead to neuronal damage and hindlimb hypersensitivity (Suter et al., 2007; Detloff et al., 2008; Grace et al., 2014; Watson et al., 2014). Here, no dLAN-elicited differences in thermal or mechanical thresholds were observed at the final timepoint, which may imply that dLAN had minimal effect on neuroinflammation after SCI.

Despite the established role of the circadian clock in governing inflammation within the CNS (Fonken et al., 2015, 2016b, 2016a; Brancaccio et al., 2019; Wang et al., 2020; Guzmán-Ruiz et al., 2023), and prior research indicating that dLAN amplifies neuronal damage following ischemic events (Fonken et al., 2019; Weil et al., 2020), our investigation revealed that housing C57BL/6J mice in dLAN had modest effect on post-SCI symptoms and recovery outcomes.

Previous research from our group demonstrated in a rat model of contusion SCI that behavioral, physiological, and molecular rhythms are disrupted post-surgery (Gaudet et al., 2018, 2019) suggesting that circadian dysregulation may already contribute to the inflammatory response provoked by SCI. If that is the case, the circadian disruption intrinsic to SCI could mask the effects of more subtle, environmentally induced disruptions such as dLAN. Similarly, the intensity of SCI-driven neuroinflammation may simply overshadow the effects of dLAN. In either case, interventions that limit SCI-induced circadian disruption or mitigate the downstream effects of that disruption may still prove useful for improving post-SCI symptoms and functional outcomes. The experiments we describe in this study center on longer-term behavioral and recovery outcomes following SCI; one future direction is to explore the molecular responses to injury and exposure to dLAN. Because the most salient changes in the SCI-elicited immune response occur acutely after injury (Ahuja et al., 2017; Gaudet and Fonken, 2018), future studies could include acute to long-term timepoints to assess alterations in circadian clock function and clarify potential mechanisms underlying long-term recovery and behavioral outcomes.

Alternatively, the effects of dLAN on neuroinflammation may be strain dependent. We used the common C57BL/6J mouse strain for this study while prior work on the effect of dLAN on ischemic stroke used Smith Webster mice (Fonken et al., 2019; Weil et al., 2020). Notably, the C57BL/6J strain, like many common inbred strains, does not produce melatonin (Roseboom et al., 1998; Kennaway, 2019) while there is evidence that the outbred Smith Webster strain does (Estrada-Reyes et al., 2018). Melatonin, along with glucocorticoids, is a primary endocrine clock output and is involved in synchronizing peripheral rhythms (Kennaway and Wright, 2002; Meléndez-Fernández et al., 2023) and dLAN reduces nocturnal melatonin secretion in rats (Molcan et al., 2019; Rumanova et al., 2020). Although melatonin has diverse and complex interactions depending on the specific physiological and experimental context, it seems to largely have an anti-inflammatory effect on microglia (Hardeland, 2018, 2021), suggesting that it would have a neuroprotective role in CNS injury models. Assuming Smith Webster mice secrete physiologically-relevant levels of melatonin in a light-sensitive manner, this could explain the limited dLAN effect we saw in this study relative to prior ischemic models (Fonken et al., 2019; Weil et al., 2020), as well as differences in lesion size and locomotor recovery observed between Swiss Webster and C57BL/6J mice after complete transection SCI (Noristani et al., 2018). Further research involving confirmed melatonin-secreting strains—e.g., CBA and C3H (Ebihara et al., 1986; Kennaway, 2019)—or direct administration of melatonin could help clarify this issue (Genovese et al., 2005).

Finally, we note that in this study and much of the prior literature, the effects of dLAN are examined in nocturnal mice. Although the molecular clock is functionally and mechanistically similar between diurnal and nocturnal species (Challet, 2007), some downstream effects vary between the two (Kalsbeek et al., 2008; Shuboni et al., 2012; Fonken and Nelson, 2014) making this an important translational consideration. Notably, nocturnal mice and diurnal Nile grass rats both exhibit enhanced immune responses when housed in dLAN, which is a key similarity that has implications for neuroinflammation (Fonken et al., 2012, 2013b; Mendoza, 2021).

### Conclusions

Our study revealed that dLAN after SCI in C57BL/6J mice slightly increased locomotor recovery at 28 dpo, exacerbated SCI-driven mechanical hypersensitivity at 13 dpo, and reduced juvenile social exploration (a measure of anxiety- and depressive-like behavior). Overall, the effect of post-SCI dLAN on locomotor deficits, pain-like behaviors, and mood-related behaviors was less far-reaching than we had predicted. Although dLAN had modest behavioral affects and did not exacerbate neurotoxicity, future studies should further address whether circadian disruption after SCI and other neurologic disorders exacerbates outcomes. Exploring circadian disruption using other manipulations – such as sleep fragmentation, stress, and lighting; alone or in combination – could more effectively model the ICU experience. Studying circadian disruption in melatonin-competent or diurnal strains and species might unmask a more human-like circadian response to injury. Conversely, given that SCI perturbs circadian rhythms (Gaudet et al., 2018, 2019), future research will reveal whether expediting reinstatement of daily rhythms could enhance recovery. Ultimately, underexplored links between the circadian system and SCI- relevant processes could aid development of new therapeutic strategies that enhance recovery after SCI.

### Funding and Disclosure

*Conflict of interest:* The authors declare no competing financial interests.

## Acknowledgements

We sincerely appreciate the guidance and support of our Lived Experience consultants, who meet with us monthly to share knowledge related to the human experience after spinal cord injury. We thank the Animal Resources Center (ARC) husbandry staff at the Health Discovery Building for excellent animal care. Partial support was provided by University of Texas at Austin start-up funds (ADG), the Wings for Life Foundation (ADG), and Mission Connect, a program of the TIRR Foundation (ADG). Research reported in this publication was supported by the National Institute for Aging under Award Number R01AG078758 (LKF) and the National Institute Of Neurological Disorders And Stroke of the National Institutes of Health under Award Number R01NS131806 (ADG). The content is solely the responsibility of the authors and does not necessarily represent the official views of the National Institutes of Health.

---

**Supplemental Figure 1.**
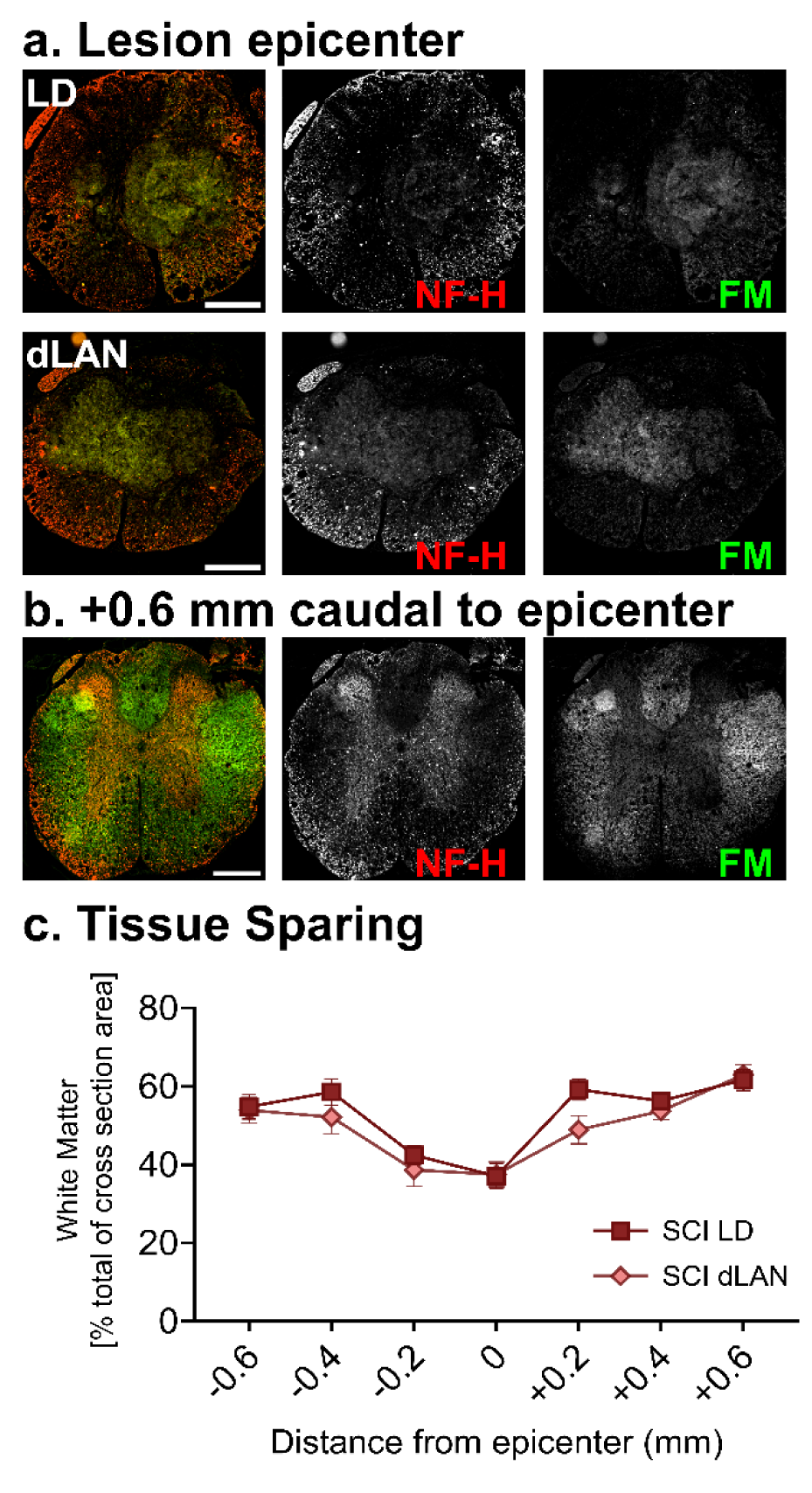
dLAN did not significantly affect post-SCI white matter sparing in C57BL/6J mice. 10 µm spinal cord cryosections from 35 dpo mice were imaged at regular 0.2 mm intervals up to -0.6 mm rostral and +0.6 mm caudal of the lesion epicenter. **A.** Representative epicenter spinal cord cross-sections from SCI mice exposed to LD or dLAN. Immunohistochemistry was performed using anti-Neurofilament H (NF-H) and FluoroMyelin Green (FM) to detect axons and myelin respectively. **B.** Largely intact white matter from a section located 0.6 mm caudal to the epicenter is shown for reference. **C.** Spared tissue was determined by tracing the intact NF-H+ FM+ area—leaving out debris within the lesion—and is expressed as a percentage of the total cross-sectional area. Lighting condition had no significant effect on white matter sparing based on two-way repeated measures ANOVA (*F_1,14_* = 2.35, *p* = 0.148). Plots show mean ± SEM. Scale bars represent 300 µm.

## Notes

https://odc-sci.org/data/956

### Competing Interest Statement

The authors have declared no competing interest.

### Summary of Updates

Improved justification of research design and methods. Enhanced discussion of results. Additional results related to white matter sparing after spinal cord injury and light-at-night.

