## Supplemental Fig for "Effects of dim light at night in C57BL/6J mice on recovery after spinal cord injury"

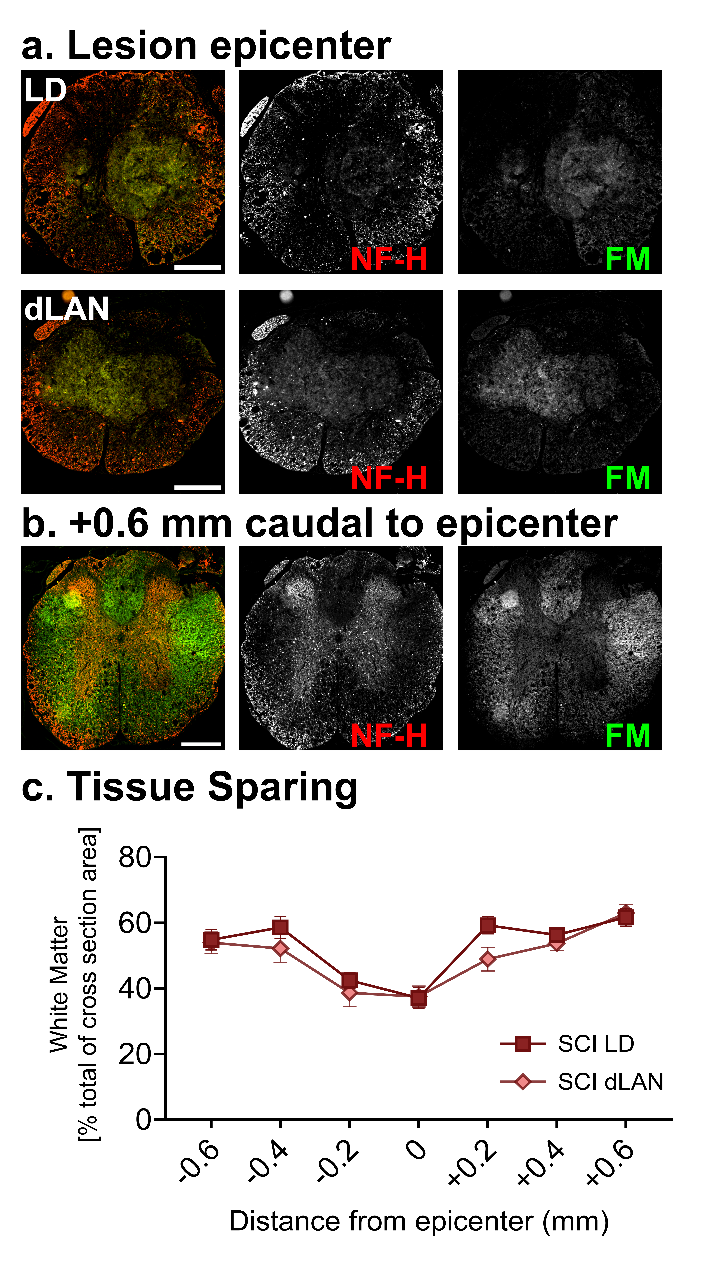

**Supplemental Figure 1**. **dLAN did not significantly affect post-SCI white matter sparing in C57BL/6J mice.** 10 µm spinal cord cryosections from 35 dpo mice were imaged at regular 0.2 mm intervals up to -0.6 mm rostral and +0.6 mm caudal of the lesion epicenter. **A.** Representative epicenter spinal cord cross-sections from SCI mice exposed to LD or dLAN. Immunohistochemistry was performed using anti-Neurofilament H (NF-H) and FluoroMyelin Green (FM) to detect axons and myelin respectively. **B.** Largely intact white matter from a section located 0.6 mm caudal to the epicenter is shown for reference. **C.** Spared tissue was determined by tracing the intact NF-H+ FM+ area—leaving out debris within the lesion—and is expressed as a percentage of the total cross-sectional area. Lighting condition had no significant effect on white matter sparing based on two-way repeated measures ANOVA (*F_1,14_* = 2.35, *p* = 0.148). Plots show mean ± SEM. Scale bars represent 300 µm.
